# Meffil: efficient normalisation and analysis of very large DNA methylation samples

**DOI:** 10.1101/125963

**Authors:** Josine Min, Gibran Hemani, George Davey Smith, Caroline Relton, Matthew Suderman

**Author notes:** Corresponding Author: Matthew Suderman. These authors equally contributed to this work (Josine Min, Gibran Hemani).

## Abstract

**Background:** Technological advances in high throughput DNA methylation microarrays have allowed dramatic growth of a new branch of epigenetic epidemiology. DNA methylation datasets are growing ever larger in terms of the number of samples profiled, the extent of genome coverage, and the number of studies being meta-analysed. Novel computational solutions are required to efficiently handle these data.

**Methods:** We have developed *meffil*, an R package designed to quality control, normalize and perform epigenome-wide association studies (EWAS) efficiently on large samples of Illumina Infinium HumanMethylation450 and MethylationEPIC BeadChip microarrays. We tested *meffil* by applying it to 6000 450k microarrays generated from blood collected for two different datasets, Accessible Resource for Integrative Epigenomic Studies (ARIES) and The Genetics of Overweight Young Adults (GOYA) study.

**Results:** A complete reimplementation of functional normalization minimizes computational memory requirements to 5% of that required by other R packages, without increasing running time. Incorporating fixed and random effects alongside functional normalization, and automated estimation of functional normalisation parameters reduces technical variation in DNA methylation levels, thus reducing false positive associations and improving power. We also demonstrate that the ability to normalize datasets distributed across physically different locations without sharing any biologically-based individual-level data may reduce heterogeneity in meta-analyses of epigenome-wide association studies. However, we show that when batch is perfectly confounded with cases and controls functional normalization is unable to prevent spurious associations.

**Conclusions:** *meffil* is available online (https://github.com/perishky/meffil/) along with tutorials covering typical use cases.

## Introduction

DNA methylation is the addition of methyl groups to cytosine bases in the DNA sequence, most often in the context of a CpG dinucleotide cytosine followed by a guanine. The addition or loss of methyl groups is often associated with changes in gene expression, and through epigenome wide associations studies (EWAS) it has been shown to associate with a wide range of complex traits. A number of technologies have been developed for interrogating DNA methylation including microarrays and sequencing-based methods. The Illumina Infinium HumanMethylation450 BeadChip (450k array) can be used to measure DNA methylation of 485k CpG sites, comprising just under 2% of the total genomic CpG content and mainly clustered around the transcription start sites of genes [1]. The new Illumina Infinium MethylationEPIC BeadChip (EPIC array) expands this coverage to ˜850k sites to include enhancer regions identified by ENCODE [2] and FANTOM5 [3].

Batch effects present a well-known challenge to microarray analysis, particularly in datasets composed of thousands of samples since they cannot all possibly be processed at the same times and by the same technical personnel. This unwanted variation can increase both false negative and false positive rates, and controlling for this is not trivial, especially as sample sizes continue to grow.

Following the popularity of quantile normalization for analyzing gene expression microarrays [4], many variations based on quantile normalization have been developed for DNA methylation microarrays (e.g. [5-7]), but they require that global methylation does not vary between samples [8]. When this does not hold, most notably between tumor and normal samples, between different tissue types, or when there are batch differences between cases and controls, quantile normalization can remove biological variation along with technical variation (e.g. [9, 10]). A feature of 450k and EPIC arrays is the inclusion of control probes - probes which do not assay biological variation and only vary due to technical effects. Functional normalization (FN) [9] exploits control probes to separate biological variation from technical variation, and its performance evaluated alongside other approaches is often favourable (e.g. [7, 9-12]).

Many DNA methylation datasets have now been generated independently, and associations between CpGs and exposures, complex traits and disease risk continue to be discovered. This rapid growth in data is warranted to improve statistical power to detect associations, however, it comes with a number of challenges that have not been fully resolved. First, existing software tools for normalising DNA methylation levels cannot be used for datasets comprising thousands of samples. Second, sharing of individual-level data is prohibited due to ethical considerations, and so the use of meta-analysis is liable to introduce heterogeneity when datasets are normalised under different procedures. Other challenges include identifying low quality methylation measurements, discovering and rectifying sample mismatches, harmonizing datasets containing both 450k and EPIC array data, removing confounding effects of cell type heterogeneity, and assessing the quality of observed associations.

With all of these challenges in mind, we have developed *meffil* to provide solutions in a user-friendly and open source R package (https://github.com/perishky/meffil). Figure 1 shows the *meffil* work-flow from raw data to quality control to normalized data to EWAS. In this paper we describe its implementation and evaluate the computational and statistical advantages that it achieves, while demonstrating where limitations might still exist.

**Figure 1.**
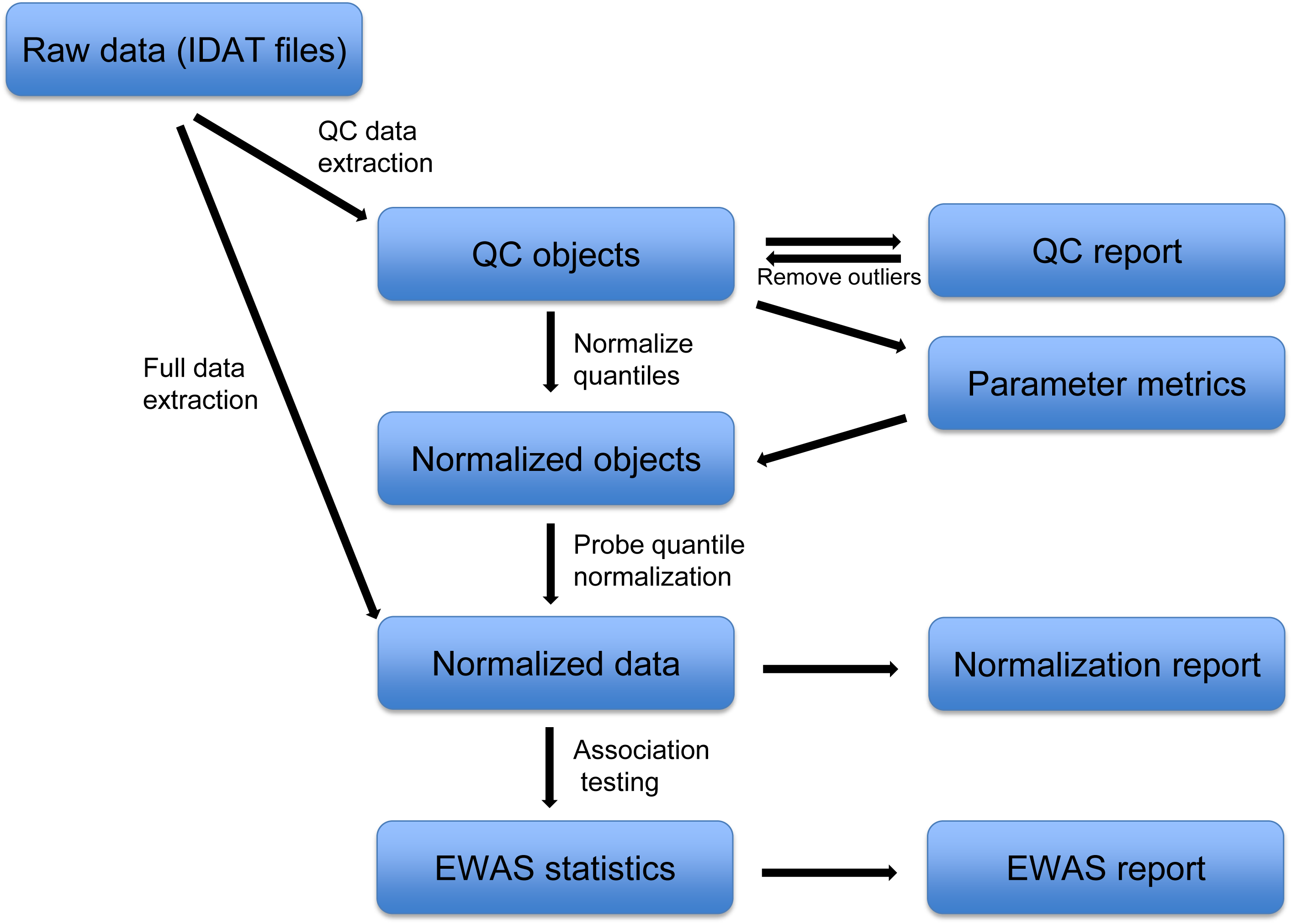
The workflow of meffil.

## Results

### Automated normalisation for heterogeneous data with improved computational efficiency

#### Computational efficiency

Our original motivation for creating *meffil* was an inability to successfully normalize ˜5400 450k arrays using available software tools. The main impediment was the large memory requirement of loading all data into memory before normalization could be initiated. We discovered, however, that functional normalization [9] could be reimplemented in a way that uses a small fraction (˜1/20) of the memory required by the entire dataset. In particular, we realized that functional normalization could be completed one sample at a time while holding in memory a relatively small summary of probe intensities for each sample. The summary consists of a control probe matrix and probe intensity quantiles. After the summary has been collected, functional normalization then proceeds to normalize intensity quantiles by removing control probe variation. Normalized methylation levels for each sample can then be derived from the normalized quantiles independently of all other samples.

To minimize running time, the *meffil* implementation makes use of the R *parallel* package to allow normalization of multiple samples simultaneously. Normalization of ˜5400 450k arrays took 3 hours on a compute server with 64 Gb of RAM and 16 processors. The memory requirements to normalize the same dataset using other software tools were too large to complete on this server. Table 2 compares the memory requirements of *meffil* and *minfi*, the most popular R package for normalizing methylation data. Most other popular packages [13, 14] that provide functional normalization capability are simply wrappers for the *minfi* implementation.

#### Scalable pipeline and reporting mechanisms

Normalization and analysis of datasets, particularly large datasets, is rarely automatic. Most of the time, problems or unexpected results are encountered requiring interactive problem-solving. Ideally, then, analysis tools should reflect this, allowing for some level of automation while allowing high-level tasks to be broken down into more specific tasks with customizable solutions. Although it would be most convenient for users to follow and interact with analyses using graphical user interface packages [15, 16], such interfaces are often not available on computational servers, particularly when the servers are nodes in a high performance computing cluster. In *meffil* we address these challenges by providing functions that nearly completely automate the entire process but can be replaced with calls to a sets of functions that allow more detailed interaction with data processing. After each main processing step (quality control, normalization and EWAS), HTML reports are generated that summarize the results of each (Figures S1-3), allowing the user to evaluate the success of each step before proceeding to the next. We also provide extensively tested QC protocols on the meffil wiki website (https://github.com/perishky/meffil/wiki).

#### Analysis of mixed 450k and EPIC datasets

Given the large number of datasets that have 450k DNA methylation profiles and the apparent popularity of the new EPIC microarray, it will likely be necessary to merge 450k and EPIC datasets for analysis. This is made possible in *meffil* by applying identical methods to probes common to both microarrays. We have yet to assess the performance of this approach due to the lack of an available mixed dataset. Fortin, et al. [17] have made a first attempt using the *minfi* package but their assessment dataset includes only three EPIC microarrays supplied by the manufacturer.

### Extending functional normalisation to reduce technical variation

Upon experimentation with functional normalization, we discovered some weaknesses and inconvenient features. We therefore sought to incorporate improvements. We assessed the performance of proposed changes by using *meffil* to process original raw data from the Accessible Resource for Integrative Epigenomic Studies (ARIES, http://www.ariesepigenomics.org.uk/) [18] comprising 4,854 methylation measurements in whole blood sampled at three time points in the life course of study participants and two timepoints in the life course of their mothers, all members of the Avon Longitudinal Study of Parents and Children (ALSPAC). Although the utmost care was taken in the generation of the high quality methylation profiles in ARIES, practical constraints lead to inconsistencies in the way samples were collected and processed. For example, DNA was extracted from a variety different sample types: whole blood, white cells, peripheral blood lymphocytes and blood spots, each with slight differences in the resulting methylation measurements. We exploit this heterogeneity to evaluate the performance of functional normalization. An EWAS of prenatal tobacco exposure was then applied to the cord blood samples comprising white cells and bloodspots (n=861). Performance was assessed by comparing resulting association statistics in ARIES to 5,801 associations of a large EWAS meta-analysis of prenatal smoking in 6,685 samples [19] (see Methods). Our EWAS of prenatal tobacco exposure in ARIES is representative of most published EWAS of large numbers of human participants. A prenatal tobacco exposure EWAS is also representative in that it identifies associations with a variety of effect sizes and significance levels, allowing us to assess the sensitivity and specificity of different analysis pipelines for identifying both strong and weak associations. As there are multiple options for selecting covariates to include in the EWAS regression model, we simplified the analysis by including only surrogate variables as covariates obtained by applying Independent Surrogate Variable Analysis (ISVA) [20]. As previously shown, these appear to sufficiently account for confounding factors including cell count heterogeneity [21].

We note that EWAS in *meffil* actually fits four different regression models (no covariates and user-supplied covariates with or without surrogate variables obtained by applying ISVA or SVA) and compares the results in an EWAS report (see Methods for more details).

#### Extending functional normalization to include fixed and random effects

As noted above, functional normalization uses control probe summaries to identify and to remove technical variation in the data. Due to microarray design, all technical variation must be captured by 42 control probe summaries, a number that will likely be too small for some of the large datasets that are currently being generated. In addition, we and others [22] have found that functional normalization often fails to completely remove slide effects (Table 2). We therefore revised our implementation of functional normalization to allow additional fixed and random effects to be included with the control matrix. Again, we used this implementation to compare EWAS performance of prenatal tobacco exposure against the associations of a large meta-analysis [19]. After explicitly including slide as a random effect, we observed increased specificity and sensitivity to detect known associations (Figure 2). Area under the curve (AUC) increases from 0.628 to 0.653 (p < 2.2 × 10^-16^, DeLong’s test).

**Figure 2.**
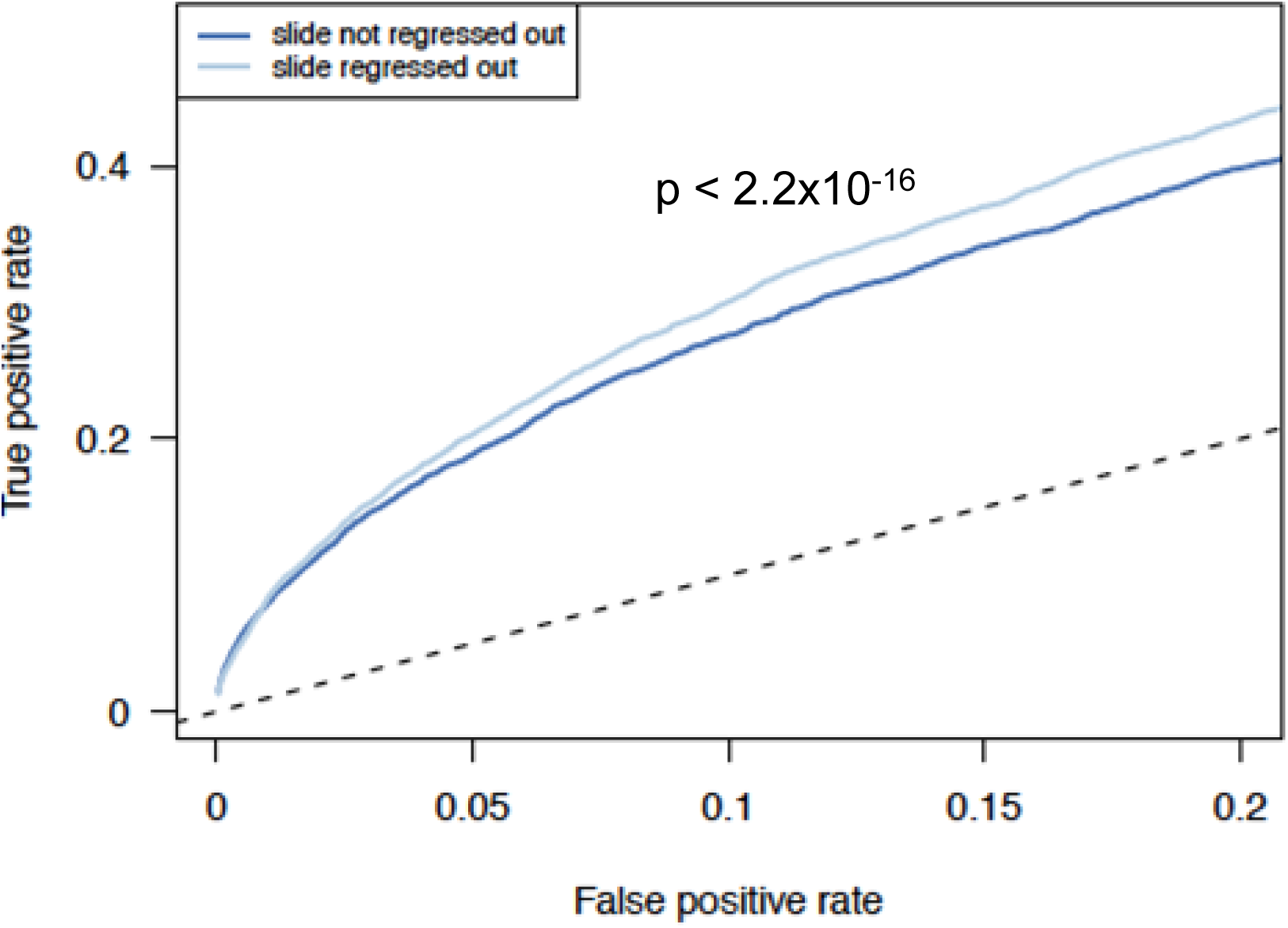
Effect of including ‘slide’ as a random effect. True positive rates are consistently higher in a downstream EWAS when variation due to ‘slide’ is removed from probe quantiles along with control variation in functional normalization. Analysis was performed in ARIES cord blood samples and associations with prenatal tobacco exposure tested. Surrogate variables obtained by applying ISVA were included as EWAS covariates. True positive rates were estimated by comparison to associations from a large meta-analysis {Joubert, 2016 #3}. Area under the curve (AUC) when slide is regressed out is larger than when it is not (AUC = 0.653 vs 0.628, p < 2.2 × 10-16, DeLong’s test).

#### Automated parameter selection

Functional normalization has one main parameter that can be set by the user: the number of principal components of the control matrix to be used to normalize the probe quantiles [9]. The maximum number is 42, the number of features in the control matrix. The default number advised by Fortin et al. [9] is two, derived as the number maximizing discovery of differentially methylated signals in a few examples. They do describe, however, other examples for which large numbers are optimal. To remove uncertainty when presented with novel data, we implemented an approach that estimates the number of principal components as the number that best explains variation in the probe intensity quantiles. This test is performed under cross validation in order to avoid overfitting. Figure 3a visualizes the unexplained variation in the probe intensity quantiles for the ARIES dataset, demonstrating a clear improvement in reducing unexplained variation at 10 principal components.

**Figure 3.**
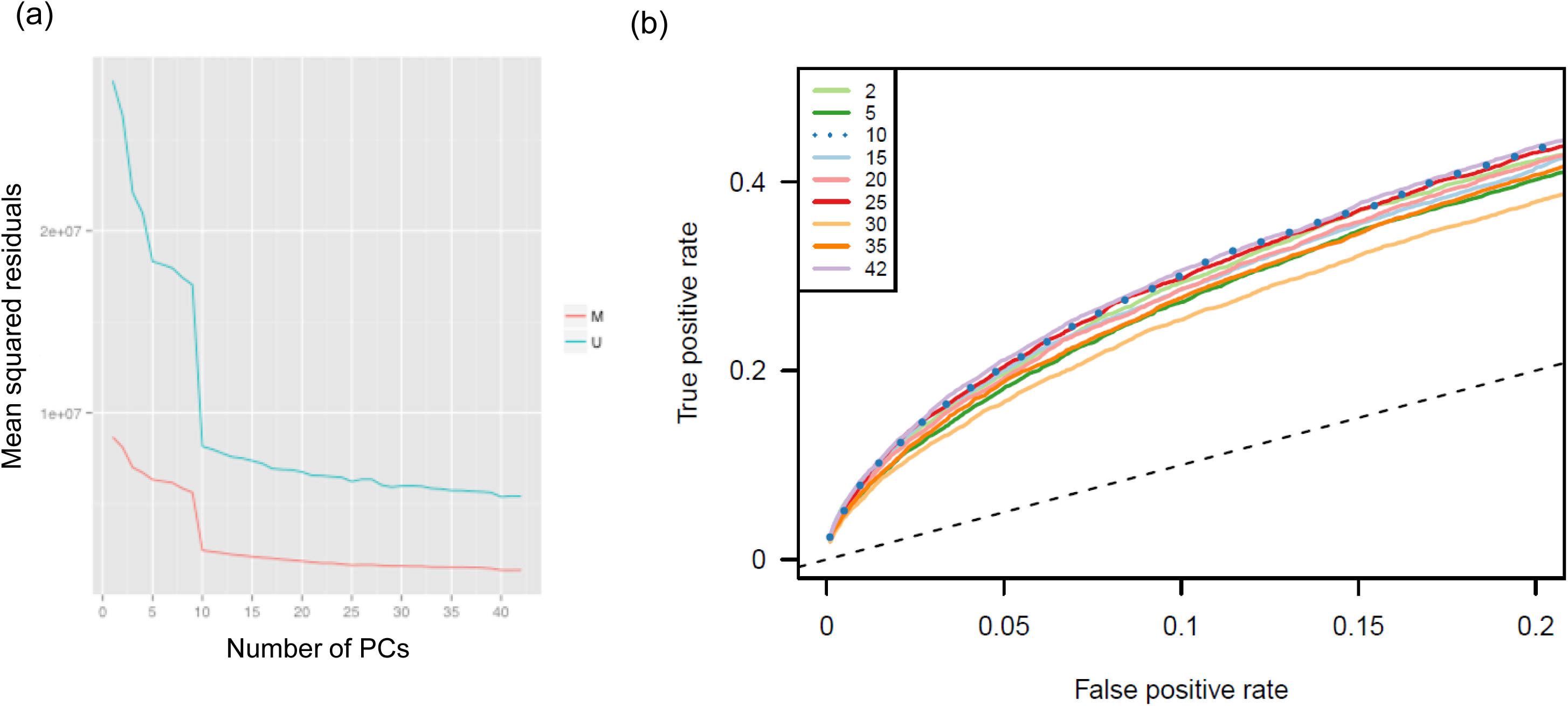
Parameter selection for functional normalization. The main parameter for functional normalization is the number of principal components of control variation with which to normalize probe quantiles. Screeplot (a) shows the metric used to meffil for choosing the optimal number of principal components, the amount of probe quantile variation unexplained by the principal components under 10-fold cross validation. Plot (b) compares true and false positive rates in a downstream EWAS of prenatal smoking in the ARIES dataset after normalizing with different numbers of principal components and regressing out slide as a random effect. True positive rates were estimated by comparison to associations from a large meta-analysis {Joubert, 2016 #3}. Both plots indicate that 10 is a reasonable choice.

To evaluate the performance of the automatic parameter selection we generated nine normalizations of ARIES cord blood samples, each normalized with a different number of control matrix principal components, and evaluated the sensitivity and specificity of identifying associations with prenatal tobacco exposure. Receiver operating characteristic (ROC) curves show that parameter choice can have a large influence (Figure 3b), with the recommended choice of 10 performing well and 42 principal components returning the best performance.

#### Reducing heterogeneity in meta-analyses with minimal data sharing

Due to the way that functional normalization is reimplemented in *meffil,* it is possible to normalize datasets residing on distinct servers together while sharing only the control matrix and probe intensity quantiles between the two servers (Figure 4a). We evaluated the effect of this approach on heterogeneity in an EWAS meta-analysis of the Accessible Resource for Integrative Epigenomic Studies (ARIES) and The Genetics of Overweight Young Adults (GOYA) study. In one meta-analysis, ARIES and GOYA were normalized separately prior to EWAS in each. In the second meta-analysis, we normalized ARIES and GOYA together prior to EWAS in each. We observed a decrease in meta-analysis heterogeneity statistics for the latter option, that is when datasets were normalized together (Figure 4b; I^2^ p = 0.047, QE p = 0.058, tau^2^ p = 0.045, H^2^ p = 0.023, Wilcoxon rank sum test of heterogeneity statistics for the top 100 associations from each meta-analysis, 134 CpG sites in total).

**Figure 4.**
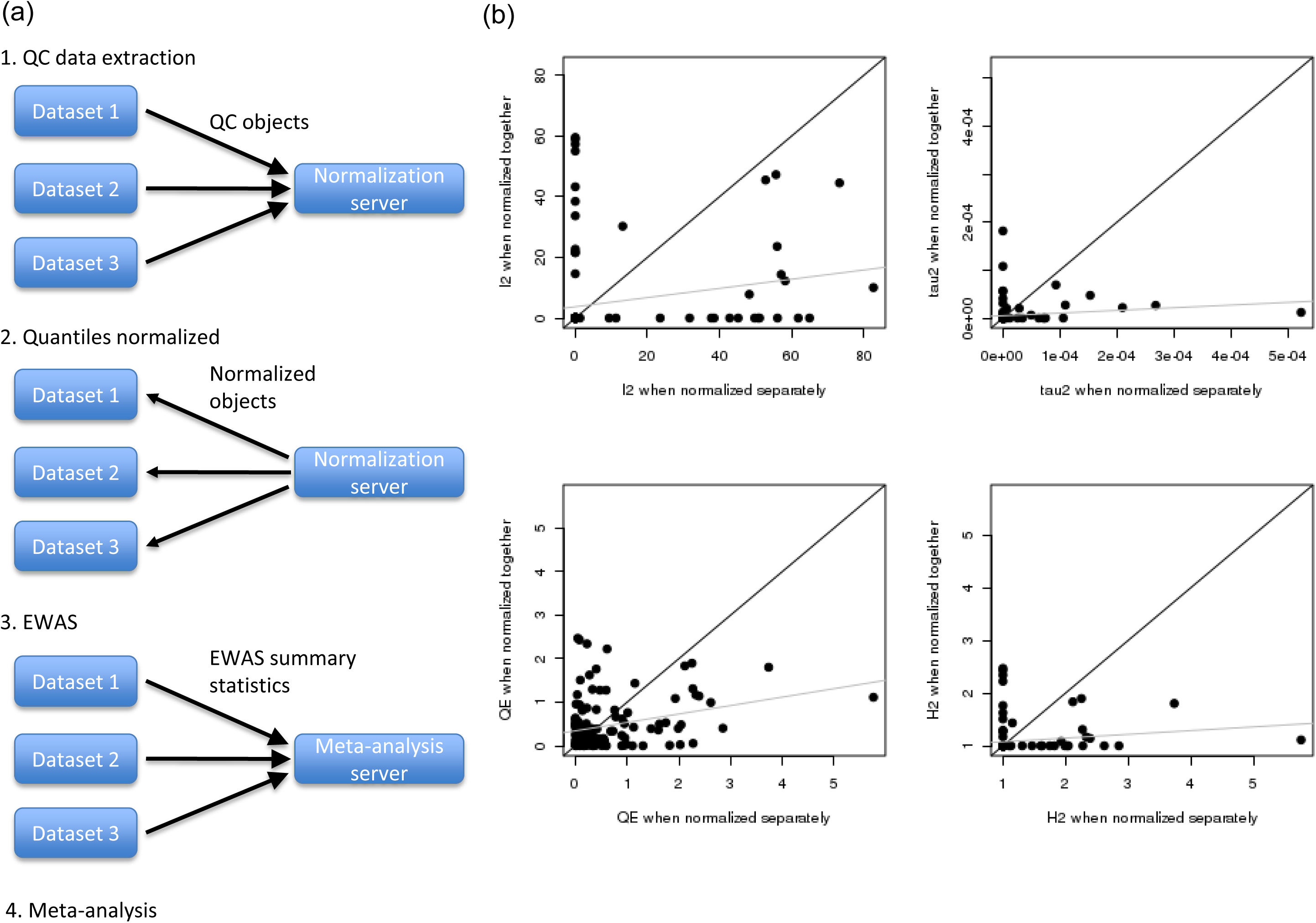
Meta-analysis with normalized data. Data can be normalized using meffil as illustrated in (a) by generating QC objects for each dataset, sending them to a normalization server for normalization and then sending them back to each dataset to complete normalization of each sample. Heterogeneity statistics are shown for meta-analyses of prenatal tobacco exposure performed on ARIES and GOYA cord samples with and without normalising the two datasets together prior to meta-analysis. Only the top 100 associations from each meta-analysis are shown comprising 134 total CpG sites. The dark diagonal line shows y=x and the gray line the regression line.

#### Perfect confounding between batch effects and biological phenotypes is not resolved by functional normalization

A common problem in epidemiological datasets is perfect confounding with batch, particularly for opportunistic case-control studies in which data is generated for cases subsequent to data collected from a control population. We evaluated the efficacy of functional normalization to remove only technical variation based on control variation while leaving biological variation intact. To test this, we compared methylation differences between methylation profiles obtained from cord blood against peripheral blood collected in adolescence under two scenarios, one in which there was perfect confounding with batch, and another in which batch was randomised across different tissues. In the unconfounded analysis, only 14 DNA methylation differences were identified (Bonferroni adjusted p < 0.05) after adjusting for cell count heterogeneity. By contrast, in the confounded analysis, there were 38950 methylation differences and this included only 7 of the 14 differences from the unconfounded analysis. Of the 38950, 62% had effect sizes in the same direction as in the unconfounded analysis. This suggests that the vast majority of the 38950 were false positives.

We then asked if adjusting for controls directly in the EWAS regression model would reduce the number of apparent false positives while retaining some of the true positives. Under this model we obtained 199 differentially methylated CpG sites, of which 50 overlapped with the 38950 from the confounded analysis and none with the unconfounded analysis. Of the 199, 123 (62%) agreed on the direction of association. Once again, these results suggest that most or all of the 199 were false positives. This was not due to the control probes failing to fully account for batch variation as a few of the ‘hybridization’ controls perfectly differentiated between batches. The false positives were then possibly due to model instability due to high correlation between controls and the variable of interest.

## Discussion

Illumina Infinium DNA methylation microarrays have been used in a number of large-scale epigenetic epidemiological studies due to their low cost and large coverage of the genome. Despite the extensive use of these arrays, memory efficient and comprehensive software are currently lacking. We have designed *meffil* to perform preprocessing, quality control, data harmonization, normalization and EWAS easily, flexibly and memory-efficiently. We have demonstrated that meffil can correct remove unwanted variation both using functional normalization and by including surrogate variables along with user variables as EWAS covariates. Automatic generation of comprehensive reports at each step allows users to assess the success of each and potentially repeat steps after tweaking parameters to improve performance.

To evaluate different settings in meffil, we used the ARIES and GOYA datasets and compared associations with prenatal tobacco exposure under various normalization schemes against those published for a large meta-analysis [19] as an example. A limitation of this approach is that the meta-analyzed set of associations might be contaminated with false positives due to batch and confounding effects that replicate across meta-analyzed datasets. Although the meta-analysis appears to be well-powered and therefore able to identify associations with small effect sizes, there are undoubtedly false negatives due to the variety of different data generation, quality control and normalization procedures applied to meta-analyzed datasets. Furthermore, all studies relied on self-reported smoking during pregnancy raising the very likely possibility of misreporting and possibly biased reporting.

We used prenatal smoking where multiple loci with small effect sizes contribute to the phenotypic variance rather than large case control effects (such as cancer). As batch effects will have the largest impact on such small effects, correcting for these effects in the most optimal way will improve power. In addition, integration and harmonization across different studies will lead to increased power in EWAS. However, simulations with different sizes of batch, confounder, and case control effects are required to find out which method and settings work best but are not the scope of this paper. Especially, as for most traits the genomic architecture is unknown, different assumptions should be made for different traits.

We and others [22] have noted that functional normalization may fail to completely remove certain technical effects, either because that variation is missing from microarray controls or because probe quantiles rather than probe intensities are directly adjusted. To address the former possibility, we allow the user to include additional technical variables as fixed or random effects. As shown in Table 1, the addition of a random ‘slide’ effect does indeed reduce variation associated with ‘slide’. For this reason, it might be better in some cases to employ a different normalization method. Crucially, we demonstrate that though functional normalization attempts to separate technical from biological variation, when batch and phenotype are perfectly confounded results can be extremely unreliable. We recommend that cases and controls be assayed jointly within a single experiment in order to obtain any value from these studies.

**Table 1.** Comparison between Minfi and Meffil applied to 1000 microarrays on a server with 16 processors and 64Gb memory.

|  | Minfi | Meffil |
| --- | --- | --- |
| Size of summary object | 2.8Gb | 194Mb |
| Memory to normalize | 0.15 | 0.05 |
| Time | 54 min | 16 min |
| Size of normalization object | 6.8G | 3.5G |

We plan in future to provide alternative normalization approaches, reimplemented in order to preserve the current low memory requirements of *meffil* and ability to normalize datasets present on distinct servers. We note however that for some methods, the reimplementation will not produce identical results because they depend on the entire dataset being loaded into memory (e.g. [7]). Future directions also include the possibility of integrating *meffil* within systems like DataShield [23] that will allow not only combined normalization but also EWAS of datasets present on distinct servers. This will improve both the power of and the speed at which meta-analyses of multiple cohort studies can be completed.

## Methods

### Data

#### Accessible Resource for Integrative Epigenomic Studies (ARIES)

Samples were drawn from the Avon Longitudinal Study of Parents and Children (ALSPAC) [24, 25]. Blood from 1022 mother-child pairs (children at three time points and their mothers at two time points) were selected for analysis as part of the Accessible Resource for Integrative Epigenomic Studies [18] (ARIES, http://www.ariesepigenomics.org.uk/). Written informed consent has been obtained for all ALSPAC participants. Ethical approval for the study was obtained from the ALSPAC Ethics and Law Committee and the Local Research Ethics Committees. Data are available from by request from the Avon Longitudinal Study of Parents and Children Executive Committee (http://www.bristol.ac.uk/alspac/researchers/access/) for researchers who meet the criteria for access to confidential data. Please note that the study website contains details of all the data that is available through a fully searchable data dictionary (http://www.bris.ac.uk/alspac/researchers/data-access/data-dictionary/).

Following DNA extraction, samples were bisulfite converted using the Zymo EZ DNA Methylation^TM^ kit (Zymo, Irvine, CA). Following conversion genome-wide methylation was measured using the Illumina HumanMethylation450 BeadChip. The arrays were scanned using an Illumina iScan, with initial quality review using GenomeStudio. During the data generation process a wide range of batch variables were recorded in a purpose-built laboratory information management system (LIMS). The LIMS also reported quality control (QC) metrics from the standard control probes on the 450k BeadChip for each sample. Samples failing QC were excluded from further analysis and the assay repeated. In total there are 5469 samples for five timepoints (birth=1127; childhood=1086; adolescence=1073; pregnancy=1100; middle aged mums=1083) measured belonging to the 1022 mother-child pairs. Sample QC and normalization was completed using with *meffil* in R version 3.2.0. Briefly, 4904/5469 ARIES samples have been successfully genotyped [26]. 112/5469 samples failed genotype QC due to sample swaps, gender mismatches, high IBD or relatedness issues between mums and kids and were removed from ARIES. We found 411 genotype mismatches (with a concordance below 80%) between the 65 snp probes on the 450k array and the genotype arrays with a concordance below 80% and these samples are removed. Furthermore, samples were removed if: i) mum samples had more than 90% concordance with a kids sample (22 samples) ii) concordance was below 80% between duplicates and less than 80% concordance with at least one other mums of kids sample (N=200). iii) mums samples of which concordance was below 80% between duplicates (N=10) iv) samples with low concordance (below 80%) with other timepoints (N=10). v) mums samples with low concordance (below 80%) with other timepoints (N=24). Methylation quality was checked by: sex check (N=191), the median intensity methylated vs unmethylated signal for all control probes (N=63), dyebias (N=14), detection pvalue (N=166), low bead numbers (N=2) and post normalization checks (N=13). Finally, 4593 samples passed QC. Samples were normalized using functional normalization using *meffil* (see below).

ARIES was normalised using 10 control probe principal components derived from the technical probes informed by *meffil* scree plots (Figure 3). These plots are not generated by other packages but shows that more than the default value of 2 PCs may be used. PCA of the normalized data using the 20,000 most variable probes shows that slide and sample type effects in cord blood were not fully eliminated by normalization (Table 2). Further investigation of these batch effects showed that slide/plate effects were confounded by sample type in this data set because, for example, different sample types were not randomized across slides and plate.

**Table 2.** Association of normalized data with ‘slide’.

|  | Association between slide and data principal component (F-statistic) |  |
| --- | --- | --- |
| Principal component | Default FN | FN with slide random effect |
| PC1 | 3.72 | 3.36 |
| PC2 | 5.53 | 4.07 |
| PC3 | 3.43 | 3.92 |
| PC4 | 13.39 | 11.20 |
| PC5 | 2.99 | 3.99 |

#### Genetics of Overweight Young Adults (GOYA)

The Genetics of Overweight Young Adults (GOYA) study is described in [27]. It includes a subset of 91,387 pregnant women recruited to the Danish National Birth Cohort during 1996–2002. Of 67,853 women who had given birth to a live born infant, had provided a blood sample during pregnancy and had BMI information available, 3.6% of these women with the largest residuals from the regression of BMI on age and parity (all entered as continuous variables) were selected for GOYA. The BMI for these 2451 women ranged from 32.6 to 64.4. From the remaining cohort, a random sample of similar size (2450) was also selected. In total, 3908 mothers were successfully genotyped. DNA methylation data were generated for the offspring of 1000 mothers in the GOYA study, equally distributed between “cases” with a BMI>32 and “controls” who were sampled from the remaining BMI distribution.

All data was imported into R version 3.2.0 and processed using *meffil*. In total there are 1010 samples belonging to 1000 children. Samples were extracted from cord blood. Ten samples were poor quality samples and were therefore repeated in the lab. 933/1010 GOYA samples have been successfully genotyped. Samples were removed due to i) genotype mismatches between 65 genotypes extracted from the genotype and 65 snp probes extracted from methylation arrays. Furthermore methylation quality was checked by: sex mismatches (23 samples), the median intensity methylated vs unmethylated signal for all control probes (N=8), bisulfate 1 probes (N=8), bisulfate II probes (N=2), dyebias (N=0), detection pvalue (N=7), low bead numbers (N=0) and post normalization checks (N=5). The data was normalized using functional normalization (Fortin et al.) and 15 PCs were used to capture technical variation. We found that slide effects and slide row were large even after normalization and we adjusted for slide and row in all analyses. After cleanup, we have 957 samples including 4 replicates. GOYA was normalized using 15 control probe principal components based on *meffil* scree plots. In our analysis we included 535 samples with a normal BMI distribution.

### Implementation of functional normalization

#### Approaches to improving efficiency

*meffil* is designed around a reimplementation of functional normalization as implemented in the *minfi* R package [9]. Output using default settings and without enhancements is therefore identical to *minfi.*

*meffil* uses the *illuminaio* R package [28] to parse Illumina IDAT files into QC objects (Figure 1). These objects play a central role in all downstream normalization procedures in *meffil*. They contain raw control probe summaries, quantile distributions of raw probe intensities, poor quality probes based on detection p-values and number of beads, predicted sex, predicted cell counts when a cell type reference is specified, and batch variable values. As in functional normalization, control probes are summarized as 42 different control types which are organized as a control matrix with one row for each control type and one column for each sample.

Quality control and normalization is made memory efficient by retaining only this small summary of the IDAT for each sample. Each is at least ˜20x smaller than the complete data. This summary object is all that is needed to perform quality control, sample and CpG site filtering, identification of batch effects, and the normalization of sample quantiles, the first normalization step of functional normalization. In this step, probe intensity quantiles are normalized between samples by fitting linear models with these quantiles to the top principal components of the control matrix. The resulting quantile residuals for each QC object are retained as a set of normalized quantiles. The normalized quantiles are then used in the second normalization step where the raw probe intensities for each sample are adjusted to conform to its set of normalized quantiles. Thus, at this point, each sample is normalized independently of all other samples.

This memory-reducing innovation makes it possible to perform the second normalization step on small subsets of the dataset, each at different times or on different compute servers. Obviously parallelization of the normalization is possible when either a single compute server has multiple processors or the normalization is being performed on a compute cluster. After the second normalization step has been completed for each individual sample, the resulting normalized methylation data subsets may be merged into a single dataset for DNA methylation analyses. The order or server on which the samples were normalized does not affect the final normalized values in any way.

#### Quality control features

*meffil* includes several features for identifying and addressing problems in the data. Quality control reports can be generated in order to uncover variation due to technical artefacts, identify outliers and flag poor quality probes and samples using detection p-values, number of beads, ratio of unmethylated/methylated signal, dye-bias and control probe checks. The report also provides checks for sample swap detection using SNP discordance between methylation and GWAS array SNP data as well as a gender check (Figure S1).

To assess the quality of a normalization, *meffil* generates a report comparing the strength of associations between batch variables with control probes and with normalized data. ANOVA and posthoc tests are used to identify problematic batches, e.g. a specific slide with technical artefacts that are not sufficiently resolved by normalization. The normalization report visualises these results with coefficient plots and a table with ANOVA F and posthoc t-statistics that pass a user-defined significance threshold (Figure S2).

All reports are generated in markdown and HTML. It is possible to convert markdown (http://daringfireball.net/projects/markdown/) output files to wide variety of other formats using tools such as pandoc (http://pandoc.org/).

#### Automatic selection of normalization parameters

Different to the implementation in *minfi*, *meffil* provides two new features for improved outputs. First, in the original implementation of functional normalization, the number of principal components of the control matrix to be included in the normalization is left as a user-defined parameter and is set to a default value of 2. *meffil* provides a method to identify the number of principal components that minimizes the residual variance unexplained by the given number of principal components. Residual variance is calculated under a 10-fold cross-validation scheme in order to avoid overfitting (Figure 3).

#### Adjusting for measured batch effects

A second feature was introduced because we observed that functional normalization failed to completely remove the variance due to certain technical artefacts such as sample slide or slide row. To address this, we allow the user to normalize sample quantiles using additional fixed and random effects. Random effects are handled using the *lme4* R package [29].

#### EWAS implementations

To deliver a comprehensive and integrated toolkit for methylation analysis, *meffil* also provides a epigenome-wide association study (EWAS) pipeline. Confounding effects are handled by including appropriate covariates in the EWAS, either as known entities or as unknown and obtained by surrogate variable analysis [20, 30, 31]. In order to better understand how each type of covariate is influencing model fitting, *meffil* in fact fits four different regression models: no covariates, only supplied covariates, supplied and surrogate variables obtained by SVA [30, 31], and supplied and surrogate variables obtained by ISVA [20]. If cell counts for samples are known, they can be included with supplied covariates. Otherwise, *meffil* allows estimation of cell counts from DNA methylation profiles based on several publicly available blood reference datasets including three cord blood references [32-34] and one peripheral blood reference [35]. Estimates may also be calculated from user-supplied references.

EWAS results are summarized in a report that includes quantile-quantile, Manhattan, covariate and variable-of-interest plots as well as tables and scatterplots showing the strongest as well as user-defined candidate CpG site associations. Outputs are displayed to allow comparison between each of the different EWAS models (Figure S3).

#### Analysis that protects study participant privacy

Because the control probes capture only technical variation, they are fundamentally non-disclosive. Given that datasets are functionally normalized in *meffil* one sample at a time and with only the control matrix and probe intensity quantiles for all samples in the datasets loaded in memory, it is possible to use *meffil* to normalize datasets residing on distinct servers together while sharing only the control matrix and probe intensity quantiles between the two servers (Figure 3a). Actual phenotype or DNA methylation levels need never be shared. The normalization proceeds by first generating QC objects for each sample in each dataset containing the control matrix and probe intensity quantiles and sending these to a single server to normalize the probe intensity quantiles. The normalized quantiles would then be sent back to each server and used to derive normalized methylation values within each corresponding dataset. The sharing of this small amount of control and probe quantile information should not violate most cohort participant privacy agreements because the information cannot be used to identify individuals.

We normalized 861 ARIES and 535 GOYA cord blood samples both separately and together and performed an EWAS of prenatal tobacco exposure in each dataset normalization. All regression models included surrogate variables and covariates as described below. We then meta-analyzed (inverse variance fixed effects) the associations, obtaining association statistics for each CpG site for the two datasets normalized separately and together.

## EWAS of prenatal tobacco exposure in ARIES and GOYA

Before analysis, samples were removed if they were replicates or due to population stratification in the genotype data. We then used three iterations to remove methylation values that were 10 SD from the mean. Associations of maternal tobacco exposure were tested in cord blood DNA methylation. In all EWAS, surrogate variables obtained by applying Independent Surrogate Variable Analysis were included as covariates [20].

### Performance evaluation

To evaluate the performance of normalization parameters and statistical methods, an appropriate gold standard for analysis output is required. Ideally, the gold standard would be a set of EWAS associations discovered in a well powered dataset and replicated in independent data. A close approximation is the 6,073 associations of prenatal tobacco exposure in cord blood DNA methylation discovered in a recent meta-analysis of 13 birth cohorts [19], one of the largest EWAS studies to date. It is not, however, a perfect standard (which may not exist) due to different normalization and analysis methods used for each dataset and lack of replication testing. To ensure that the associations were not contaminated with false positives due to genetic and technical artefacts, we retained the 5,801 of the 6,073 not linked to probes potentially affected by genetic variants (MAF > 0.01 in the European subset of 1000 Genomes) or prone to non-specific binding [36].

We evaluated performance by constructing Receiver Operating Characteristic (ROC) curves using the truth data described above. We used the CRAN ROCR package [37] and code provided by [12] to construct the ROC curves. For all ISVA analyses we set the seed to 160815 (15 August 2016). Meta analyses were conducted using an inverse variance fixed effects model using the metafor R package [38].

## Acknowledgements

We are extremely grateful to all the families who took part in the ALSPAC study, the midwives for their help in recruiting them, and the whole ALSPAC team, which includes interviewers, computer and laboratory technicians, clerical workers, research scientists, volunteers, managers, receptionists and nurses.

The UK Medical Research Council and Wellcome (www.wellcome.ac.uk; 102215/2/13/2) and the University of Bristol provide core support for ALSPAC. This publication is the work of the authors and Matthew Suderman will serve as guarantor for the contents of this paper. This research was specifically funded by the UK Economic and Social Research Council (www.esrc.ac.uk; ES/N000498/1 to CR); all authors work within the Medical Research Council Integrative Epidemiology Unit at the University of Bristol, which is supported by the UK Medical Research Council (www.mrc.ac.uk; MC_UU_12013/1 and MC_UU_12013/2).

---

**Figure S1.**
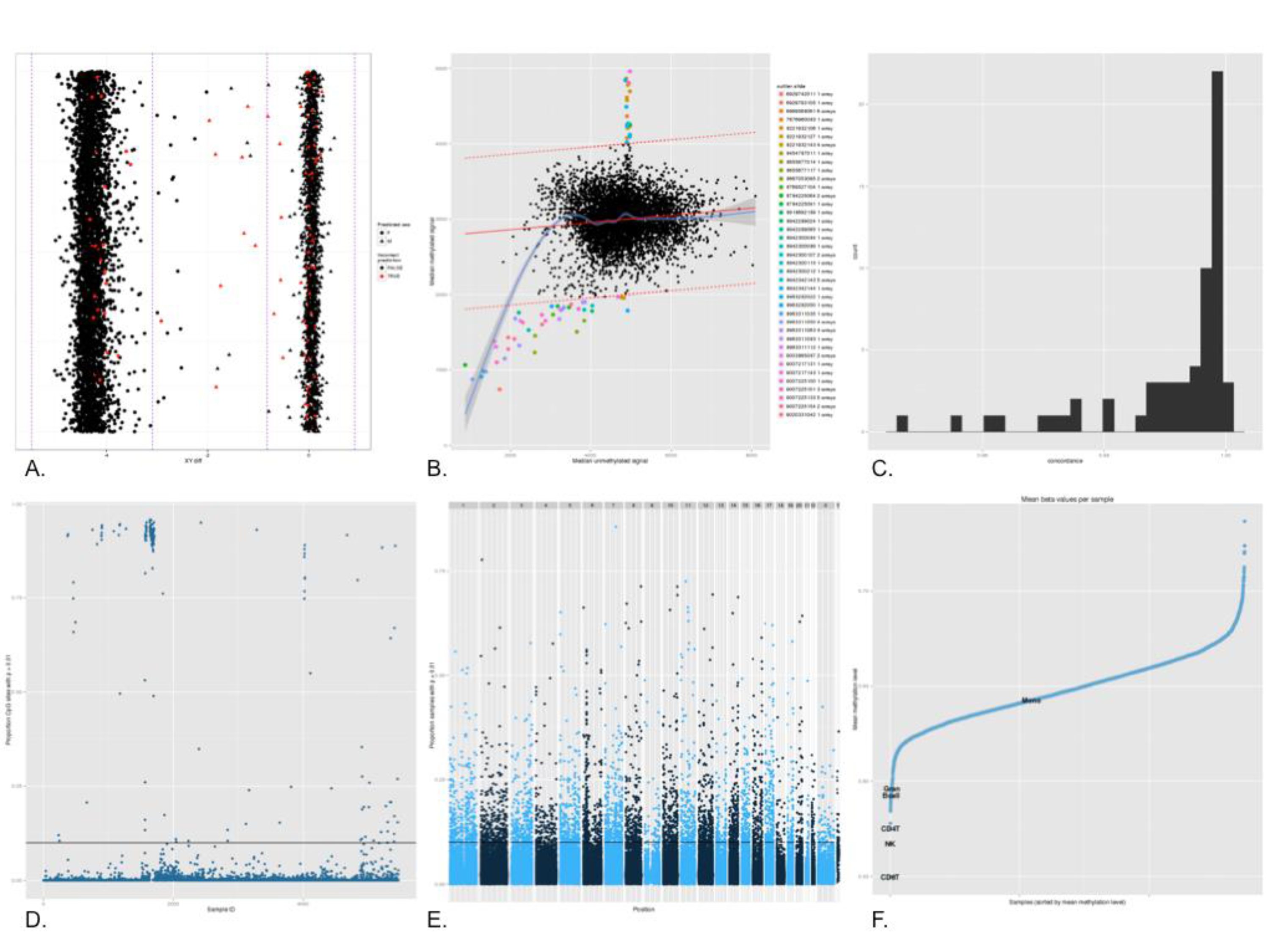
meffil QC report. Plots generated in the QC report. Results are shown for 5469 samples from the ARIES resource. A. Gender prediction. B. Comparison of methylated versus unmethylated signal C. GWA discordance using 65 SNP probes D. Proportion of detected probes by sample. E. Proportion of detected samples by probe. F. Methylation levels used to estimate cell counts for each sample versus reference methylation profiles.

**Figure S2.**
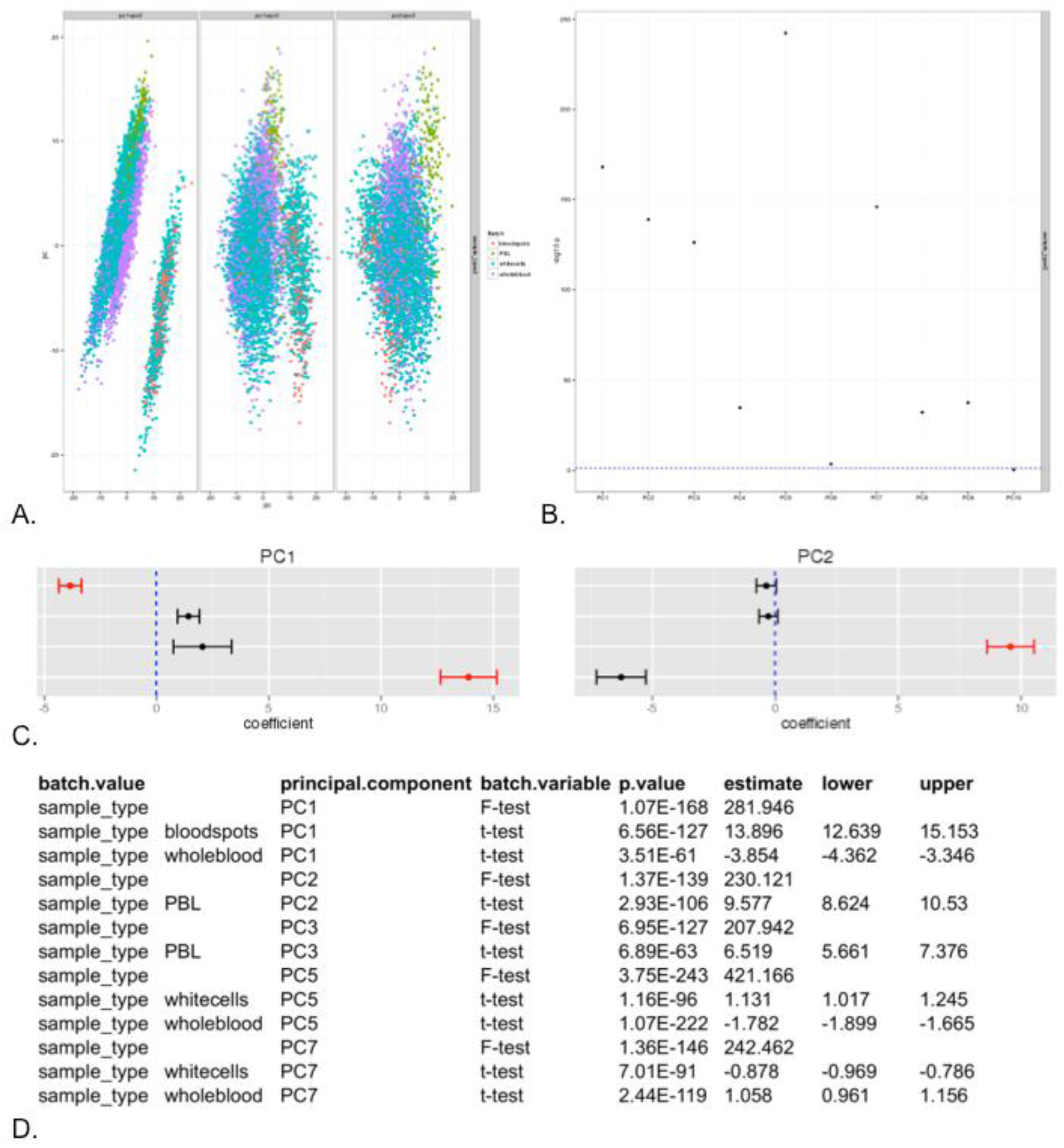
meffil normalisation report. Plots generated in the normalization report. A. PCA plot colored by batch variable. B. Anova test pvalues between PCs extracted from the normalized betas and batch variable. C. Coefficient plots for associations between PCs and a batch variable.

**Figure S3.**
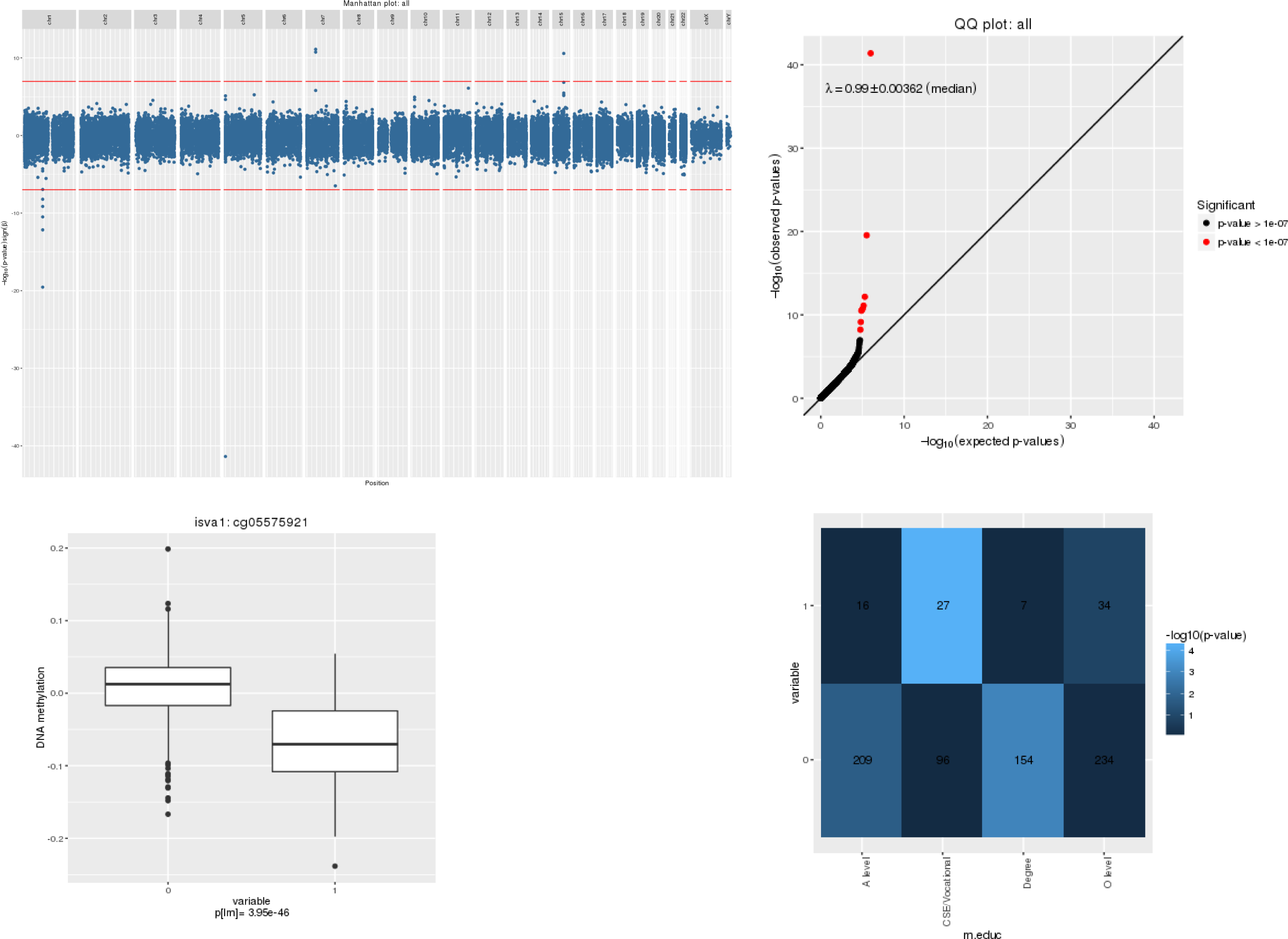
meffil EWAS report. Plots generated in the EWAS report. Results are shown for an EWAS on prenatal smoking in ARIES (777 cord samples). A. Manhattan plot. B. QQ plot C. Methylation differences between cases and controls for a CpG of interest. D. Association between confounder and phenotype (variable of interest).

